# Progenitor Timing Shapes NG2-Glia Fate and Oligodendrocyte Differentiation

**DOI:** 10.1101/2025.09.24.678191

**Authors:** Ana Cristina Ojalvo-Sanz, Carolina Pernia-Solanilla, Laura López-Mascaraque

## Abstract

The developmental identity and fate of NG2-glia remain debated: are they transient oligodendrocyte precursors or a distinct, self-renewing glial population? Here, we examined how the temporal origin of progenitors influences NG2-glia and oligodendrocyte lineages in the dorsal cortex. Using in utero and postnatal StarTrack electroporation at E12, E14, E16, and P0, we traced their progeny to P30 and P90, performing a clonal analysis in adult mouse brains.

Progenitors labeled at E16 generated significantly larger and more widely dispersed NG2-glia clones, whose contribution increased from P30 to P90, suggesting enhanced proliferative capacity. In contrast, P0-derived progenitors showed reduced NG2-glia maintenance and a strong bias toward oligodendrocyte differentiation, forming larger OL clones. Clonal heterogeneity, including mixed NG2-glia/OL clones, was observed across all stages but peaked at E16.

These results identify E16 as a critical window for NG2-glia expansion and self-renewal, while P0 marks a transition toward oligodendrocyte lineage restriction, establishing a developmental framework for adult NG2-glia heterogeneity and maybe a regenerative potential.

**Significance Statement:** We show that developmental timing and progenitor identity shape NG2-glia and oligodendrocyte fates in the postnatal cortex. NG2 progenitors at E16 generate expansive, widely dispersed NG2-glia clones, whereas P0 progenitors bias toward oligodendrocyte differentiation, revealing critical temporal windows that establish long-term glial heterogeneity.

## INTRODUCTION

The development of the central nervous system (CNS) depends on the precisely orchestrated generation of both neurons and glial cells from neural progenitor cells (NPCs) (Casingal et al., 2022; Li et al., 2021; Noctor et al., 2008). Within the glial lineage, NG2-glia occupy a particularly intriguing niche. Traditionally classified as oligodendrocyte precursor cells (OPCs) (Nishiyama et al., 2009; Trotter et al., 2010), these cells have long been viewed as transient intermediates en route to forming mature, myelinating oligodendrocytes. However, accumulating evidence challenges this classical perspective (Bedner et al., 2020; Bergles et al., 2000; Boda & Buffo, 2014; Dansu et al., 2021; Kamen et al., n.d.; Marques et al., 2016; Minocha et al., 2015; Miralles et al., 2023; Parolisi & Boda, 2018; Steinhäuser & Dietrich, 2017; Viganò et al., 2016).

A significant fraction of NG2-glia appears to resist terminal differentiation, persisting throughout life as a stable, self-renewing population with diverse and specialized functions. These include supporting neuronal migration during development, forming dynamic synaptic-like contacts with neurons, and interacting with neurons and pericytes to influence vascular physiology (Bergles et al., 2000; Hesp et al., 2018; Lepiemme et al., 2022; Steinhäuser & Dietrich, 2017; Yalçın & Monje, 2021). Such observations challenge the traditional view of NG2-glia as transient progenitors and raise fundamental questions about the mechanisms governing their fate. Are NG2-glia merely stalled progenitors, or do they represent a distinct, lineage-stable glial type shaped by developmental timing and environmental context?

Addressing this question requires a better understanding of when and how NPCs commit to either an oligodendrocyte fate or a persistent NG2-glia identity. This issue is essential to delineate glial lineage progression and to uncover the origins of glial heterogeneity. While previous studies have examined NPC proliferation and differentiation at discrete developmental stages (García-Marqués et al., 2014; Marques et al., 2016; Miralles et al., 2023), a systematic and comparative analysis across embryonic and perinatal time points has remained elusive. It is still unclear whether early, multipotent progenitors generate NG2-glia that persist into adulthood or whether these cells gradually transition into the oligodendrocyte lineage over time. Moreover, the contributions of later, more fate-restricted progenitors to the balance between NG2-glia maintenance and oligodendrocyte differentiation are poorly understood.

In this study, we address these unresolved questions by investigating how the developmental timing of NPC labeling influences the fate of NG2-glia and oligodendrocytes in the postnatal cortex. Using in utero electroporation with StarTrack-based lineage tracing vectors, we labeled NPCs at distinct embryonic and perinatal stages (E12, E14, E16, and P0) and analyzed their progeny at postnatal day P30 and P90. By distinguishing clones derived from GFAP-expressing versus NG2-expressing progenitors, we systematically compared the contributions of different progenitor pools to NG2-glia and oligodendrocyte lineages over time.

Our clonal analyses reveal that both progenitor identity and the temporal window of labeling influence whether NG2-glia persist or differentiate. Notably, progenitors labeled at E16 gave rise to a larger proportion of NG2-glia that expanded between P30 and P90, whereas those labeled at P0 generated fewer NG2-glia and showed a stronger bias toward oligodendrocyte differentiation. These findings highlight the existence of critical temporal windows during which NG2-glia fate is differentially tuned—either to preserve the progenitor pool or to fuel oligodendrogenesis.

By dissecting the developmental dynamics that govern NG2-glia identity, this study fills an important gap in our understanding of glial lineage specification. Beyond their developmental significance, these insights have translational implications: clarifying how NG2-glia balance self-renewal and differentiation may inform strategies for enhancing remyelination, designing cell-based therapies, and promoting repair in demyelinating disorders of the CNS. To our knowledge, this is the first systematic clonal analysis directly comparing NG2-glia fate across multiple embryonic and perinatal time points, providing a novel framework for understanding their lineage dynamics.

## RESULTS

### Temporal Windows Define NG2-Glia Versus Oligodendrocyte Fate

To understand how the developmental timing of neural progenitor cell (NPC) labeling shapes the lineage output of their progeny—either maintaining an NG2-glia identity or differentiating into oligodendrocyte precursor cells (OPCs)—we performed *in utero* electroporation using the StarTrack system at embryonic days E12, E14, and E16, as well as postnatal day 0 (P0). This strategy enabled long-term clonal tracing of NPCs and their progeny, which were analyzed at postnatal days P30 and P90. Cell types were identified based on morphological features and marker expression (Figure 1A). NG2-glia exhibited small soma, highly branched processes (Figure 1B), and PDGFRα immunoreactivity, whereas mature oligodendrocytes were identified by their rounded nuclei, aligned myelin processes, and expression of the APC marker (Figure 1C).

**Figure 1.**
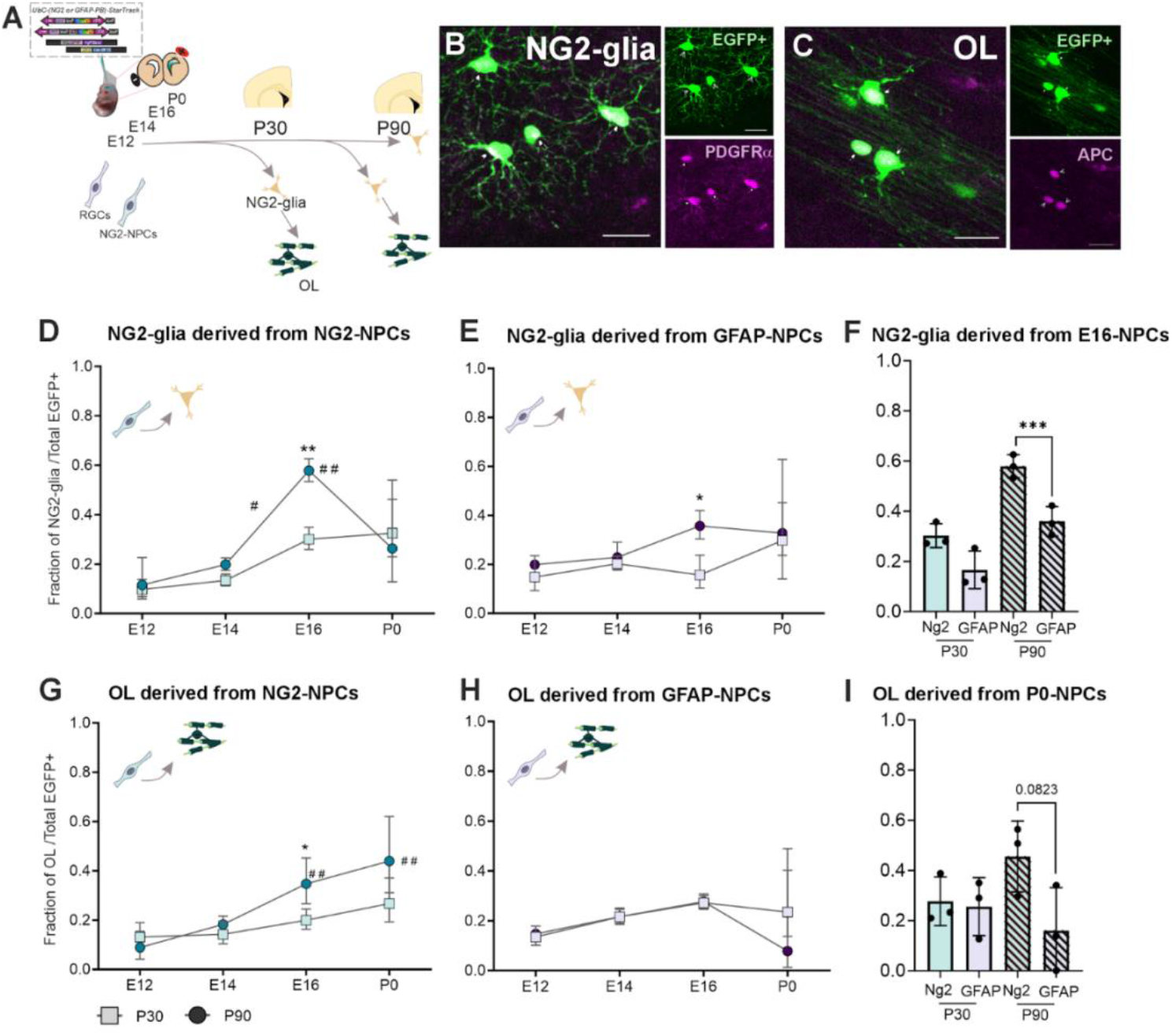
Developmental timing and progenitor identity influence NG2-glia and oligodendrocyte fate. (A) Experimental design. Neural progenitor cells were labeled using in utero or postnatal electroporation of the StarTrack system at E12, E14, E16, or P0. The glial progeny NG2-glia and oligodendrocytes (OL) were analyzed at P30 and P90 to assess lineage fate. (B) Representative image of NG2-glia identified as PDGFRα^+^ cells. (C) Representative image of oligodendrocytes identified as APC^+^ cells. (D–E) Quantification of the proportion of NG2-glia among total EGFP^+^ cells derived from NG2-NPCs (D) or GFAP-NPCs (E) across developmental stages and postnatal time points. Squares indicated P30-derived progeny and circles P90-derived progeny. (F) Comparison of NG2-glia contribution from NG2-NPCs and GFAP-NPCs labeled at E16, analysed at P30 and P90. (G–H) Quantification of oligodendrocyte (OL) contribution from NG2-NPCs (G) or GFAP-NPCs (H). (I) Comparison of P30 and P90 OL contribution from NG2-NPCs and GFAP-NPCs labeled at P0. All data are presented as mean ± SD. Statistical comparisons were performed using two-way ANOVA followed by Bonferroni post hoc test. * p < 0.05, **p < 0.01, ***p < 0.001, for diferences between P30 and P90 derived progeny and #p < 0.05, ##p < 0.01 for within-stage comparisons Scale bar: 25um. N = 6 animals per stage (3 from NG2-NPCs and 3 from GFAP-NPCs).

To determine whether lineage fate decisions are influenced not only by developmental timing but also by progenitor identity, we selectively labeled GFAP-expressing and NG2-expressing NPCs and compared their progeny across developmental stages. Both NG2-glia and oligodendrocytes were observed in the labeled lineages. However, their relative proportions varied markedly depending on labeling time and progenitor type (Figures 1D– I).

In GFAP-NPC progeny, no significant differences in NG2-glia contribution were detected at P30 across time points (p = 0.3522). By P90, however, a significant divergence emerged (p = 0.0372; Figure 1E), indicating that temporal factors influence long-term glial identity even when initial outputs appear comparable. In NG2-NPC progeny, the effect of developmental timing was more pronounced (Figure 1D): significant differences in NG2-glia abundance were observed at both P30 (p = 0.0041) and P90 (p = 0.0126). Labeling at E16, in particular, resulted in a striking enrichment of NG2-glia (E16 vs. E12: p = 0.0121; E16 vs. E14: p = 0.0283), with proportions rising from 13.23% at E12 and 19.90% at E14 to a peak of 57.94% at E16, before declining again to 31.59% at P0 (Figure 1D).

Notably, progeny from E16-labeled NPCs exhibited a significant increase in NG2-glia between P30 and P90, both in NG2-derived (p = 0.0019) and GFAP-derived lineages (p = 0.0243). This twofold rise suggests that E16 progenitors give rise to NG2-glia populations capable of either sustained proliferation or long-term persistence (Figures 1D–E). Across all stages, NG2-glia were consistently more abundant in NG2-NPC progeny compared with GFAP-NPC progeny (p = 0.007; Figure 1F), highlighting an intrinsic lineage bias operating in addition to temporal regulation.

In contrast, oligodendrocyte generation followed a different trajectory. Among NG2-NPC progeny, oligodendrocyte contribution increased progressively with later labeling stages (p = 0.0547), whereas GFAP-derived progeny showed no significant stage-dependent variation (p = 0.3165) (Figures 1G–H). NG2-NPCs labeled at P0 yielded the highest proportion of oligodendrocytes (35.48%), significantly higher than those labeled at E12 (10.40%, p = 0.0056) and E14 (18.41%, p = 0.0237) (Figure 1G). Furthermore, oligodendrocyte output increased between P30 and P90: for instance, progenitors labeled at E16 increased from 20.20% oligodendrocytes at P30 to 35.48% at P90 (p = 0.0484), while P0-derived progenitors rose from 27.71% to 45.60% (p = 0.1446), indicating a robust trend toward terminal differentiation at later stages.

Although no statistically significant differences were detected between P0-GFAP-NPCs and P0-NG2-NPCs in oligodendrocyte output (p = 0.0823), NG2-derived progeny consistently produced higher fractions of oligodendrocytes by P90 (Figure 1I). This suggests that NG2-expressing progenitors retain a lineage-intrinsic bias toward oligodendrogenesis, which becomes most evident when they are labeled at late developmental stages.

Together, these results reveal that E16 represents a critical temporal window during which NG2-expressing progenitors preferentially generate NG2-glia that persist and expand into adulthood, whereas P0 progenitors—particularly NG2-NPCs—undergo a shift toward oligodendrocyte differentiation. The progressive increase in oligodendrocyte output from P30 to P90 underscores that late-born progenitors remain competent for terminal differentiation over extended postnatal periods. Thus, both developmental timing and progenitor identity act in concert to bias lineage allocation, ultimately shaping the balance between NG2-glia persistence and oligodendrocyte differentiation in the postnatal cortex.

### A Key Developmental Window at E16: NG2-Glia Clones Show Enhanced Expansion

To evaluate whether the temporal origin of NG2- or GFAP-expressing progenitors influences clonal fate and proliferative potential, we performed *in utero* electroporation with a combination of twelve XFP StarTrack plasmids and the PiggyBac transposase driven by either NG2 or GFAP promoters at E12, E14, E16, or P0 (Figure 2A). We then analyzed the progeny at P30, identifying 271 gliogenic clones classified into three categories: NG2-glia-only, oligodendrocyte-only (OL), or mixed NG2+OL clones (Figure 2B–F). Among these, 143 clones originated from GFAP-driven StarTrack (GFAP-PB) and 128 from NG2-driven StarTrack (NG2-PB). Restricting our analysis to these three categories ensured consistency and comparability across conditions.

**Figure 2.**
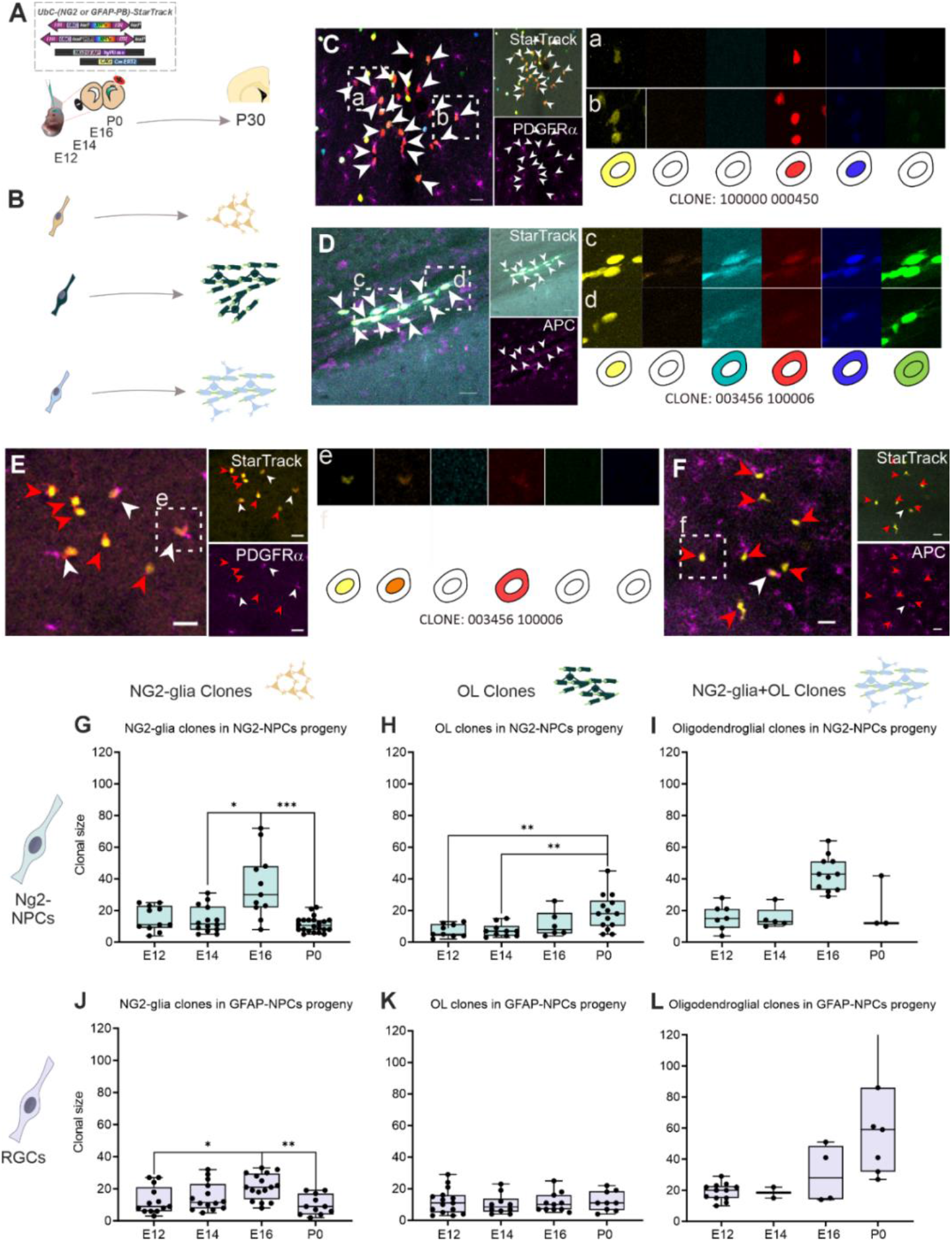
Developmental timing influences the clonal output of NG2-glia and oligodendrocytes from distinct progenitor types. (A) Schematic of in utero or postnatal electroporation of GFAP+ or NG2+ progenitors using the StarTrack genetic tool at E12, E14, E16, or P0, with analysis at P30. (B) Representation of different clone types: NG2-glia clones (orange), OL clones (dark green), and mixed NG2-glia+OL clones (blue). (C) NG2-glia clones PDGFRα+. Insets (a, b) represent cells that belong to the same clone with the color-code. (D) OL clones APC+. Insets (c, d) represent cells that belong to the same clone with the color-code. (E-F) NG2-gli+OL mixed clone with some cells PDGFRα+ (white arrows) and others APC+ (red arrows). Insets (e, f) represent cells that belong to the same clone with the color-code. (G–I) Clonal size for NG2-glia (G), OLs (H), and NG2-glia+OL clones (I) derived from NG2+ progenitors. (J–L) Quantification of clonal size for NG2-glia (J), OLs (K), and mixed clones (L) derived from GFAP+ progenitors. Data represent individual clones, with bars indicating mean ± SD. Statistical comparisons across stages were performed using one-way ANOVA with post hoc tests. *p < 0.05; **p < 0.01; ***p < 0.001. Scalebar: 50um

In the NG2 lineage, NG2-glia clones were significantly larger when labeled at E16 (median: 30 cells) compared to E12 (11 cells), E14 (11 cells), and P0 (11 cells) (Figure 2G; one-way ANOVA: p = 0.0012; E14 vs. E16: p = 0.0225; P0 vs. E16: p = 0.0005; E12 vs. E16: p = 0.0001). This represents a striking enrichment of NG2-glia expansion at E16, indicating that progenitors at this stage have a high clonal amplification capacity. In contrast, OL clones derived from NG2-progenitors remained small at embryonic stages and increased only at P0 (median: 18cellsat P0 vs. 7cells at E12 and E14; Figure 2H; P0 vs. E14: p = 0.0497), suggesting that postnatal progenitors are more prone to adopt an OPC-like fate.

Mixed NG2+OL clones appeared at all stages but were significantly larger at E16 (median: 43cells) compared to E12 (15cells), E14 (13cells), and P0 (12cells) (Figure 2I; E16 vs. E12: p = 0.0001; E16 vs. E14: p = 0.003; E16 vs. P0: p = 0.022). These results reveal that E16 NG2-progenitors are highly heterogeneous, producing diverse glial fates while sustaining robust proliferative capacity.

GFAP-derived progenitors also generated significantly larger NG2-glia clones at E16 (median: 21cells) compared to E12 (9cells), E14 (11.5cells), and P0 (9cells) (Figure 2J; ANOVA: p = 0.0042; E12 vs. E16: p = 0.0385; E16 vs. P0: p = 0.005). This indicates that even radial glia undergo a transient peak of NG2-glia production at late embryonic stages. OL clones derived from GFAP progenitors were consistently small and showed no significant differences across time points (Figure 2K), confirming that GFAP-NPCs contribute less to oligodendrocyte lineages. Interestingly, mixed GFAP-derived clones were largest at P0 (median: 59cells) compared to earlier embryonic stages (20cells at E12; 18.5cells at E14; 28cells at E16) (Figure 2L; ANOVA: p = 0.0043; P0 vs. E12: p = 0.0028), suggesting that postnatal radial glia transiently acquire NG2-like characteristics, enabling them to generate both NG2-glia and OLs.

Overall, these findings highlight E16 as a critical developmental window for clonal expansion of NG2-glia in both lineages, while the increased size of OL and mixed clones at P0 reflects a shift toward more restricted, OPC-like differentiation. The recurrent presence of mixed clones across stages underscores the heterogeneous and asynchronous nature of glial fate acquisition.

### Temporal Origin Influences Rostro-Caudal Dispersion of NG2-Glia Clones

To determine whether the developmental stage of progenitor labeling also influences the spatial distribution of gliogenic progeny, we measured the rostro-caudal (RC) dispersion of NG2-glia, OL, and mixed NG2+OL clones derived from NG2- and GFAP-expressing progenitors at P30 (Figure 3).

**Figure 3.**
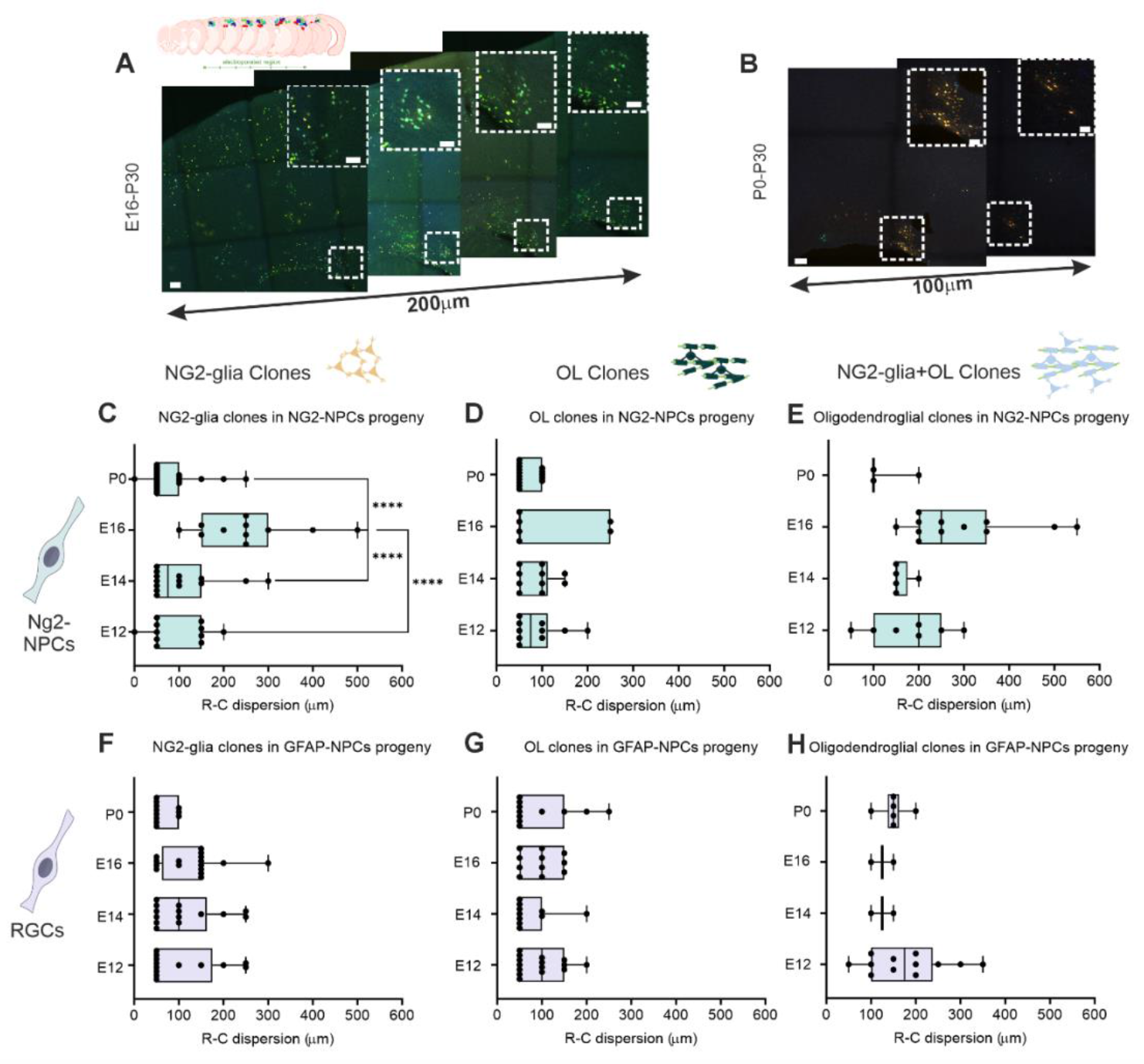
Progenitor labeling at E16 promotes broad rostrocaudal dispersion of NG2-glia clones. (A–B) Representative image of cortical sections showing the rostrocaudal (RC) dispersion of StarTrack-labeled clones at P30. (A) NG2-glia clones derived from E16-labeled NG2+progenitors exhibit broad RC distribution. (B) OL clones derived from P0 NG2+ progenitors show limited dispersion. (C–E) RC dispersion for NG2-glia clones (C), OL clones (D), and mixed clones (E) from NG2+ progenitors. (F–H) RC dispersion of NG2-glia (F), OLs (G), and mixed clones (H) derived from GFAP+ progenitors. Boxes represent interquartile ranges; whiskers, 10– 90th percentiles; dots, individual clones. Statistical significance determined by one-way ANOVA with post hoc Tukey test. ****P < 0.0001. Scalebar: 100um

In the NG2 lineage, NG2-glia clones exhibited a striking stage-dependent increase in RC dispersion, peaking at E16 (mean: 250 μm). This value was significantly higher than those observed at E12 (50 μm), E14 (75 μm), and P0 (50 μm) (Figure 3C; ANOVA p < 0.0001; E16 vs. others: p < 0.0001). These results suggest that NG2-progenitors at E16 not only expand clonally also confer on their progeny with greater migratory or proliferative capacity, enabling broader cortical coverage. In contrast, OL clones exhibited consistently limited dispersion across stages (Figure 3D), reinforcing their more spatially restricted behavior. Mixed clones displayed the broadest distributions at E16 (250 μm), compared with 200 μm at E12, 150 μm at E14, and 100 μm at P0 (Figure 3E), confirming that E16 progenitors generate highly dynamic and spatially expansive populations.

In GFAP-derived lineages, RC dispersion of NG2-glia clones did not vary with developmental stage but followed a slightly different trajectory: the widest spread was observed at E14 (100 μm), whereas E12, E16, and P0 progenitors generated clones with more limited dispersion (50 μm) (Figure 3F). OL clones from GFAP-progenitors exhibited minimal and invariant dispersion across all stages (Figure 3G). Mixed GFAP-derived clones showed relatively stable distributions, with a modest increase at P0 (Figure 3H), consistent with the idea that postnatal radial glia transiently broaden their fate and spatial influence.

Together, these findings demonstrate that E16 represents a critical window for both expansion and spatial dispersion of NG2-glia and mixed gliogenic clones, whereas P0 progenitors display a shift toward more spatially restricted, oligodendrocyte-biased outputs. Moreover, lineage-specific differences emerge clearly: NG2-derived clones consistently display broader dispersion than GFAP-derived ones, particularly at late embryonic and postnatal stages. Based on these results, we propose the model depicted in Figure 4. In this framework, early progenitors (E12–E14) in both lineages generate relatively stable ratios of NG2-glia and oligodendrocytes with limited spatial expansion. By contrast, late-embryonic NG2-progenitors (E16) undergo a burst of clonal amplification and dispersion, producing NG2-glia clones that persist and expand between P30 and P90.

**Figure 4.**
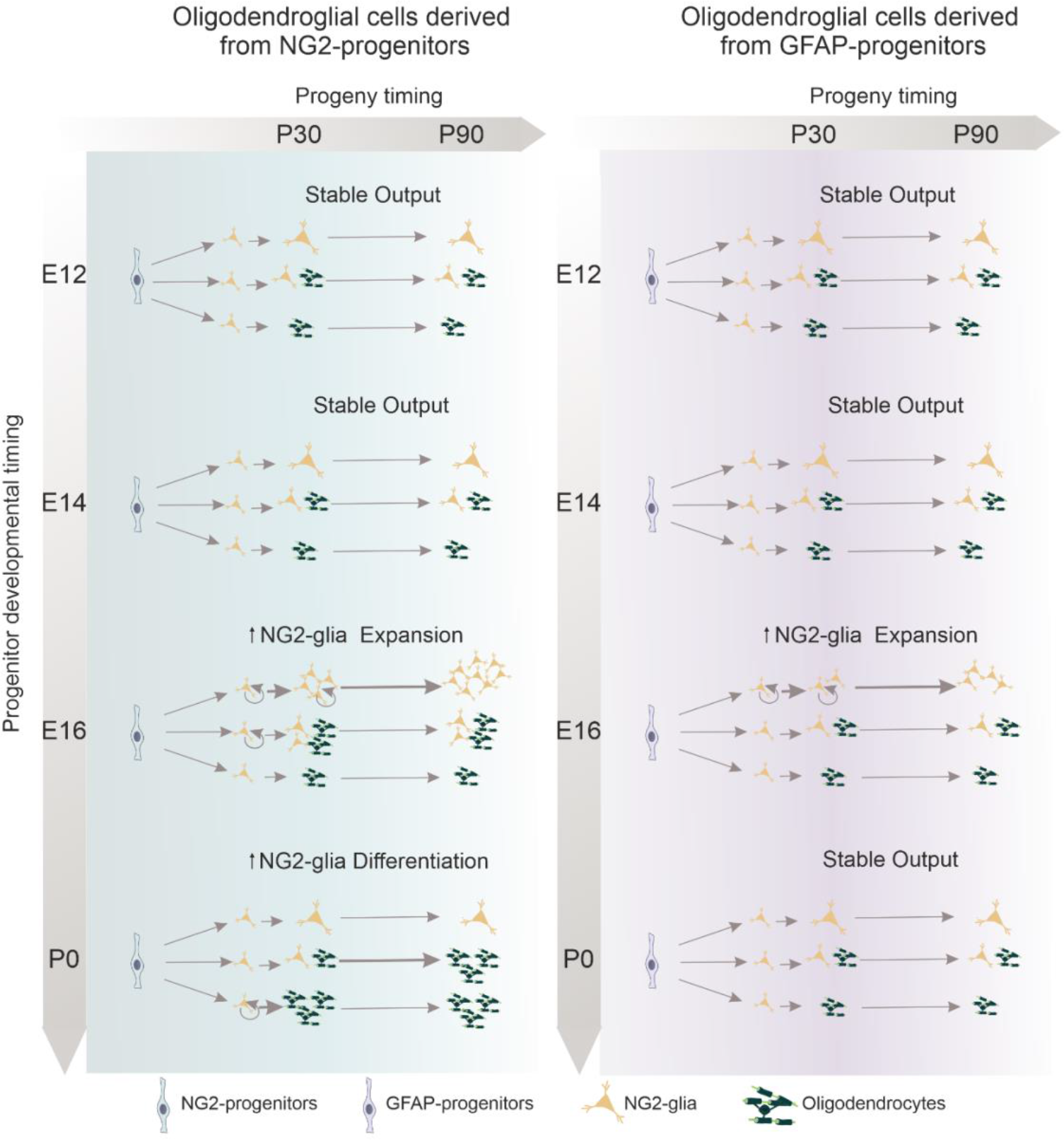
Proposed model for the temporal and lineage-dependent generation of NG2-glia and oligodendrocytes in the postnatal cortex. Schematic summary integrating clonal and progeny analyses, illustrating our working hypothesis. Neural progenitors (NG2-progenitors on the left and GFAP-progenitors on the right), labeled at different embryonic and perinatal stages (E12, E14, E16, P0), give rise to distinct proportions of NG2-glia (orange) and oligodendrocytes (dark green) at P30 and P90. Early progenitors (E12–E14) maintain stable ratios of NG2-glia and oligodendrocytes over time in both lineages. Late-embryonic progenitors (E16) generate the largest NG2-glia clones, which expand from P30 to P90, that may reflect higher proliferative activity (circular arrow). In contrast, perinatal NG2-progenitors (P0) may preferentially generate larger oligodendrocyte clones and show a marked decline in NG2-glia proportion from P30 to P90, accompanied by a corresponding increase in oligodendrocytes, potentially driven in part by the differentiation of NG2-glia within mixed NG2 and oligodendrocyte clones. However, GFAP-progenitors maintain more stable lineage outputs, similar to those of early-stage progenitors.

In contrast, perinatal NG2-progenitors (P0) shift toward OPC-like behavior, generating larger oligodendrocyte clones and displaying a pronounced decline in NG2-glia output, likely reflecting widespread differentiation of NG2-glia into oligodendrocytes within mixed clones. GFAP-derived progenitors, in contrast, exhibit more stable and restricted patterns of contribution, resembling early-stage progenitors.

## DISCUSSION

Our study reveals that both developmental timing and progenitor identity influence the generation, expansion, and fate specification of NG2-glia and oligodendrocytes (OLs) in the postnatal dorsal cortex. Through lineage tracing and clonal analysis across embryonic and early postnatal stages, we show that progenitors labeled at embryonic day 16 (E16) generate NG2-glia with enhanced proliferative and dispersive potential, whereas progenitors labeled at postnatal day 0 (P0) preferentially produce NG2-glia that subsequently undergo biased differentiation into OLs. These findings provide a framework for understanding how temporal and molecular properties of progenitors become encoded into long-term oligodendroglial behavior.

E16 emerges as a critical temporal window in oligodendroglial lineage progression. At this stage, progenitors—particularly NG2-expressing NPCs—produce significantly larger NG2-glia clones with broader rostro-caudal dispersion than those labeled at earlier embryonic (E12, E14) or later postnatal (P0) stages. This suggests that NG2-glia derived from E16 NPCs are uniquely primed for expansion and spatial distribution, likely reflecting exposure to stage-specific developmental signals. By birth, however, this proliferative program shifts toward a more fate-restricted, oligodendrogenic trajectory, consistent with previous reports that NG2-glia expand transiently before differentiating into OLs within a narrow postnatal window (Nishiyama, Serwanski, et al., 2021; Nishiyama, Shimizu, et al., 2021).

NG2-glia were long considered a homogeneous population uniformly distributed across the cortex and destined for oligodendrogenesis (Hill et al., 2011). Subsequent studies revealed important differences between gray- and white-matter NG2-glia, including proliferative capacity, electrophysiological properties, and responsiveness to neural (Dimou & Simons, 2017). More recent work has shown that NG2-glia, while initially homogeneous, diversify over time and across regions (Spitzer et al., 2019)), giving rise to functionally distinct subpopulations. Functional heterogeneity also extends to intrinsic properties such as inward K^+^ conductance, which is modulated by neuronal activity and, in turn, regulates cell-cycle dynamics and proliferation (Pivoňková et al., 2024). This aligns with the current view of NG2-glia as a plastic and heterogeneous population capable of adapting to developmental and pathological cues (Bandler et al., 2022; Boda et al., 2022). This NG2-glia diversity could be essential in disease context: in multiple sclerosis, for instance, human lesions exhibit a reduction in NG2-glia heterogeneity (Jäkel et al., 2019), while after traumatic brain or spinal cord injury, subsets of NG2-glia proliferate to support wound closure, whereas others differentiate into oligodendrocytes or astrocytes (Dean et al., 2023; Hein et al., 2025; Hesp et al., 2018). Our findings suggest that diversification trajectories imprinted during embryogenesis, particularly at E16, may preconfigure how NG2-glia respond to injury: E16-derived cells may preferentially expand to replenish lost populations, whereas P0-derived cells are biased toward differentiation to replace myelinating oligodendrocytes. Single-cell transcriptomics supports this view, revealing multiple NG2-glia states within the same developmental stage, ranging from highly proliferative to differentiation-primed populations (Bandler et al., 2022; Huang et al., 2020; Marques et al., 2016). Subpopulations marked by molecules such as GPR17 or Olig2 display different proliferative or differentiation biases, and their responses vary in disease models (Boda et al., 2011; olig2 Bonfanti et al., 2017; Miralles et al., 2023; Viganò et al., 2016). These molecular identities likely intersect with the temporal programs we identified: for example, E16 progenitors may be particularly responsive to mitogenic signals such as PDGF-AA and FGF within an extracellular matrix enriched in tenascin-C and hyaluronic acid (Baron et al., 2002),), whereas P0 progenitors may be more exposed to differentiation cues such as thyroid hormone (T3), IGF-1, or laminins (Lourenço et al., 2016; Wheeler & Fuss, 2016).

Progenitor identity also plays a decisive role. NG2-progenitors consistently produced higher proportions of NG2-glia and OLs than GFAP-progenitors at both E16 and P0, supporting an intrinsic bias of NG2^+^ NPCs toward oligodendroglial fates, potentially mediated by transcription factors such as Olig2. Indeed, Olig2 loss redirects NG2^+^ cells toward astrocytic fates, underscoring its role in lineage restriction (Nishiyama et al., 2014; Zhu et al., 2012). Our clonal analyses further revealed variability in clone size, dispersion, and lineage composition, including mixed NG2-glia/OL clones appearing at all stages. Notably, E16 NG2-derived clones exhibited the broadest rostro-caudal dispersion, highlighting a unique window of enhanced migratory and proliferative potential. While previous clonal studies have largely focused on earlier embryonic windows (García-Marqués et al., 2014; Ojalvo-Sanz & López-Mascaraque, 2021; Sánchez-González et al., 2020; Zhang et al., 2020), our data emphasize the importance of late embryonic stages in shaping long-term oligodendroglial fate. This clonal heterogeneity could be important in demyelinating conditions, where NG2-glia respond in spatially heterogeneous and clonal patterns (Boda et al., 2011; Bonfanti et al., 2017; Viganò et al., 2016).

Together, these results suggest that much of the adult heterogeneity of NG2-glia originates from developmental lineage trajectories established in embryogenesis. The identification of E16 as a pivotal stage for NG2-glia expansion and dispersion highlights the interplay between temporal programs and progenitor identity in determining the balance between persistence and differentiation. From a translational perspective, the recognition of temporally defined windows of NG2-glia competence opens new opportunities for therapeutic intervention. NG2-glia derived from late embryonic progenitors may represent a particularly valuable population for regenerative strategies, given their ability to persist, expand, and disperse. By contrast, the OL-biased progeny of perinatal progenitors may be more readily recruited for remyelination. Harnessing these differences will require a deeper understanding of the molecular cues that regulate the shift from expansion to differentiation, including extracellular matrix components, growth factors, and transcriptional programs. Future studies integrating single-cell transcriptomics, in vivo imaging, and functional assays will be crucial to link the temporal origins of NG2-glia with their regenerative potential in disease contexts. Ultimately, defining the developmental logic that shapes NG2-glia heterogeneity could inform novel cell- and stage-specific approaches to promote CNS repair in demyelinating and traumatic pathologies.

## MATERIALS AND METHODS

### Mouse Lines

Adult and pregnant wild-type C57BL/6 mice were obtained from the Cajal Institute (Madrid, Spain) and maintained under standard housing conditions (12 h light/dark cycle, 22°C) in accredited animal facilities. All procedures complied with EU Directive 2010/63/EU and Spanish regulations (RD 1201/2005, L 32/2007) and were approved by institutional and regional bioethical committees (PROEX 314/19). Embryonic day 0 (E0) was defined by the presence of a vaginal plug, whereas postnatal day 0 (P0) corresponded to the day of birth. Mice older than P30 were considered adults. For each experimental condition, a minimum of three animals per group was used. Both males and females were included in all analyses.

### StarTrack Method

Neural progenitor cells (NPCs) and their progeny were traced using StarTrack, a multicolor genetic labeling strategy based on the piggyBac transposon. This method allows stable integration of fluorescent reporters, enabling heritable and combinatorial color-coding of cell lineages (Figueres-Oñate et al., 2016; García-Marqués et al., 2014). StarTrack employs 12 plasmids encoding six different fluorescent proteins (XFPs: mT-Sapphire, mCerulean, EGFP, YFP, mKO, mCherry) driven by independent promoters. Variability in fluorophore type, subcellular localization (nuclear/cytoplasmic), and intensity yields more than 49 trillion unique combinations. For additional technical details, see (Figueres-Oñate et al., 2025; Ojalvo-Sanz et al., 2025)

The hyperactive PiggyBac transposase (hyPBase) was expressed under cell-type-specific promoters: NG2-hyPBase (provided by Prof. Sellers) to target NG2 progenitors, and GFAP-hyPBase (courtesy of Dr. Lundberg) for GFAP-lineage progenitors. Plasmid purification was performed with the NucleoBond Xtra Maxi Kit (Macherey-Nagel; #740414.50). For fate-mapping, UbC-EGFP or UbC-H2B-EGFP were used in combination with Cre-ERT2. For clonal analysis, all StarTrack plasmids were co-electroporated with Cre-ERT2 and the appropriate hyPBase.

### Electroporation Protocols

*In utero* electroporation (IUE) was performed at E12, E14, and E16. Pregnant females were anesthetized with isoflurane (3% induction, 2% maintenance) and treated with Baytril (5 mg/kg) and Meloxicam (300 µg/kg). After uterine exposure, 1–2 µL of DNA mix (1.8–2 µg/µL in distilled water, 0.05% Fast Green) was injected into the lateral ventricle (LV). Electroporation was carried out with five 50-ms pulses (950-ms intervals) using electrode tweezers (Protech International; #CUY650-P3). Voltages were adjusted by stage (E12: 33 V; E14: 35 V; E16: 37 V).

Postnatal electroporation was performed at P0 under hypothermia. Pups received 0.5 µg/µL of StarTrack DNA (with 0.2% Fast Green) into the LV. Five 100-mV pulses (50 ms) were delivered. Pups were then warmed at 37°C for recovery and allowed to develop to P30 or P90.

### Tamoxifen-Induced Removal of Episomal Copies

To activate Cre-ERT2 and eliminate episomal plasmids, tamoxifen (Sigma-Aldrich, T5648-1G; 20 mg/mL in corn oil) was administered intraperitoneally (7.5 mg/kg). Tamoxifen was administered at P8 to avoid birth complications.

### Tissue Processing and Sectioning

At designated timepoints (P30, or P90), animals were deeply anesthetized with pentobarbital (Dolethal, 40–50 mg/kg; Vétoquinol) and transcardially perfused with 4% paraformaldehyde (PFA) in 0.1 M phosphate buffer (PB). Brains were post-fixed for 1 h in 4% PFA and coronally sectioned at 50 µm using a vibratome. Serial sections were collected into six-well series for downstream analyses.

### Immunohistochemistry

Sections were washed and permeabilized in PBS-Triton X-100 (0.5%, then 0.1%). For PDGFRα staining, sections were pretreated with 10% methanol. Antigen retrieval, when required, was performed using alternating hot/cold citrate buffer cycles (3×5 min). Sections were blocked in 5% normal goat serum (NGS) in PBS-T 0.1% for ≥2 h at room temperature. Primary antibodies were applied overnight at 4°C in blocking buffer. The following antibodies were used: Sox10 (rabbit, 1:200, #69661, Cell Signaling), Olig2 (rabbit, 1:500, AB9610, Millipore), NG2 (rabbit, 1:100, AB5320, Millipore), PDGFRα (rabbit, 1:300, #31745, Cell Signaling), and APC/CC1 (mouse, 1:300, OP80, Calbiochem). Sections were subsequently incubated with Alexa Fluor-conjugated secondary antibodies (goat anti-rabbit or goat anti-mouse, Thermo Fisher) for 2 h. After washing, sections were mounted in Mowiol (Polysciences).

### Imaging Processing

Epifluorescence and confocal imaging were performed on Leica TCS-SP5 microscopes at the Cajal Institute (Madrid) and Hospital Nacional de Parapléjicos (Toledo). Fluorophore channels were acquired separately to prevent spectral bleed-through. Laser intensities ranged from 15–30%. Excitation/emission spectra: mT-Sapphire: 405/520–535 nm, mCerulean: 458/468–480 nm, EGFP: 488/498–510 nm, YFP: 514/525–535 nm, mKO: 514/560–580 nm, mCherry: 561/601–620 nm, Alexa Fluor 633/647: 633/650–760 nm. Images were processed using LAS X software (v3.5.1, Leica) to generate maximum intensity projections.

### Quantification

Quantitative analyses were performed in Fiji (ImageJ, v1.53q). One 50-µm section per 300 µm was analyzed per brain, covering the entire electroporated regions. StarTrack-EGFP+ cells were quantified and classified as NG2-glia or oligodendrocytes based on immunostaining.

### Clonal Analysis

Clones were reconstructed using LAS X and a custom Fiji macro developed by the Cajal Institute’s imaging unit. Detection thresholds for each channel were defined using fluorescence histograms, and binary masks were generated to segment individual cells. Cells sharing identical XFP combinations and subcellular localization patterns were classified as sibling clones. Morphology and marker expression were used to determine cell identity. The electroporation site was defined by the first and last sections containing labeled cells, spaced at 50 µm intervals.

### Statistical Analysis

All statistical tests were performed in RStudio (v1.4.1106) and GraphPad Prism (v6.0). Normality was assessed with the Kolmogorov–Smirnov and Dallal-Wilkinson-Lillie tests. For non-parametric comparisons, the Mann–Whitney test (two groups) or Kruskal–Wallis test (multiple groups) was used. Statistical significance was set as *p < 0.05, **p < 0.01, ***p < 0.001. Results are reported with 95% confidence intervals.

## Acknowledgments

Animal, imaging and microscopy facilities of the Instituto Cajal for the assistance.

